# Evaluation of Commercially Available Exosomal Isolation Kits from Human Plasma

**DOI:** 10.1101/808931

**Authors:** Yuzhe Sun, Hefu Zhen, Mei Guo, Jingyu Ye, Zhili Liu, Xiuqing Zhang, Yan Yang, Chao Nie

## Abstract

Exosomes are cell-derived lipid bilayer particles which are abundant in biological fluids. Exosome is an emerging source of biomarkers to diagnose various human diseases. Sequencing based exosomal studies could provide a comprehensive view of exosomal RNA and protein. To extracted these inclusions, exosomes should be isolated from the plasma first. Several exosome isolation methods were introduced since the discover of exosome. To promote the clinical application of exosomal inclusions, different isolation methods should be compared. We isolated exosomes from human plasma by using user-friendly and commercially available kits, SBI ExoQuick and QIAGEN exoRNeasy. Subsequently, small RNA sequencing was performed with two groups of isolated exosome samples and one group of plasma samples. No fundamental differences of exRNA yield between SC and EQ were found. In RNA profile analysis, the small RNA aligned reads, miRNA pattern, sample clustering varied as a result of methodological differences. Small RNA isolated by ExoQuick presented better data quality and RNA profile than exoRNeasy. This study compared sRNA sequencing data generated from two exosome isolation kits, it provides a reference for future small RNA studies and biomarker prediction in human plasma exosome.

## 1. Introduction

Extracellular vesicles (EVs) are small lipid bilayer particles that are commonly observed in human body fluids involving in cell-to-cell communication. Exosome, a major component of EVs, carries the fundamental biological elements such as protein, mRNA, long non-coding RNA (lncRNA), and, especially small RNA (sRNA) [1]. Recently, exosome has become an emerging source of biomarkers of various human diseases like neurodegenerative diseases and cardiovascular diseases[2]. Using peripheral blood to detect exosomal inclusions could satisfy the rapid and noninvasive requirement of chronic disease prediction[3]. Particularly, the exosomal sRNA including microRNA (miRNA) are of great diagnostic interest since they are important post-translational regulators and protected by lipid bilayer structure from exposure to RNases [4]. Thus, method of extracting RNA from exosomes and RNA-seq library preparation kits are developed [5-7]. Due to the nature of sRNA diversity in exosomes, exosomal sRNA patterns are easily interfered, it is extremely hard to reproduce the same results which have been obtained from the previous experiments [4]. A few works have compared different methods of exosomal sRNA extraction by measuring isolated exosome integrity and isolation efficiency from different tissues. However, previous works didn’t systematically analyze transcriptomic or proteomic data[8]. Our study focused on comparing next-generation sequencing data generated from different exosome isolation techniques.

Several techniques have been invented to isolate these small vesicles from different tissues [9]. These techniques include ultracentrifugation, density gradient separation, ultrafiltration, immunological separation and polymer-based precipitation, column-based chromatography and peptide binding [10]. Among them, ultracentrifugation is most commonly used technics, and centrifugation-based techniques are considered to be the gold standard for exosome isolation [11]. However, extremely strict experimental conditions, such as high-speed centrifuge, and 4 oC experimental environment, are required by ultracentrifugation because of the time-consuming procedures [12]. Besides, the throughput of the ultracentrifugation method is limited, while the output is unstable since isolation steps require qualified experimental skills. Therefore, ultracentrifugation approach is not suitable for the large-scale clinical trial. As exosomes in human blood undergoes a degradation-generation homeostasis, the difficulties for isolating and sequencing exocellular vesicles rely on several aspects such as instability and complexity of the samples, the collection procedures, the exosome isolation methods, the heterogeneity of physicochemical properties, the storage conditions, and low RNA content [4]. Therefore, a minimally invasive, uniformed, cost-effective exosome isolation method is necessary. Commercial kits have its own advantages. For instance, the exosome isolation procedure is easily adapted to clinical laboratory environment and the experiment circle is usually short enough so that the plasma samples can be handled on time. The stable performance of exosome isolation is also suitable for researchers in hospitals. Thus, two commercially available kits: SBI ExoQuick™ (EQ) and Spin column (SC) from QIAGEN exoRNeasy Serum/Plasma Kit, were chosen as they represent two isolation method, the polymer-based precipitation and the column-based chromatography, respectively. This study is focusing on the impact of different exosome isolation kits on sRNA sequencing data, it could contribute to the investigation of exosomes and promote the application of exosomes in the clinical practice.

## 2. Results

### 2.1 Workflow for isolating RNA from extracellular vesicles

To evaluate how the extracellular vesicle isolation method influences the small RNA profile inside vesicles, we tested two commonly used exosome isolation kits (ExoQuick™, EQ; exoEasy spin column, SC) by next-generation sequencing on the human plasma. Human plasma was used as the sample source in order to minimize the effect of tissue diversity and 46 samples of plasma were collected. Each sample was separately treated with two kits, while a blank control was set up using the same sample. Small RNA yield were measured by NanoDrop 2000 (Table 1, Figure 1). Then all samples (200ng per sample) were carried out standardized miRNA extraction, sRNA library preparation and sequence platform (Table 1). Datasets generated from three sample groups were named as EQ, SC and PC samples. Table 1 shows the schematic of the study design and the differences of three groups. Human Brain Reference RNA (HBRR) samples were added into sequence lanes as reference samples in order to eliminate the batch effect. A summary of small RNA-seq results is presented in supplementary files (Supplementary Table 1).

**Table 1.**
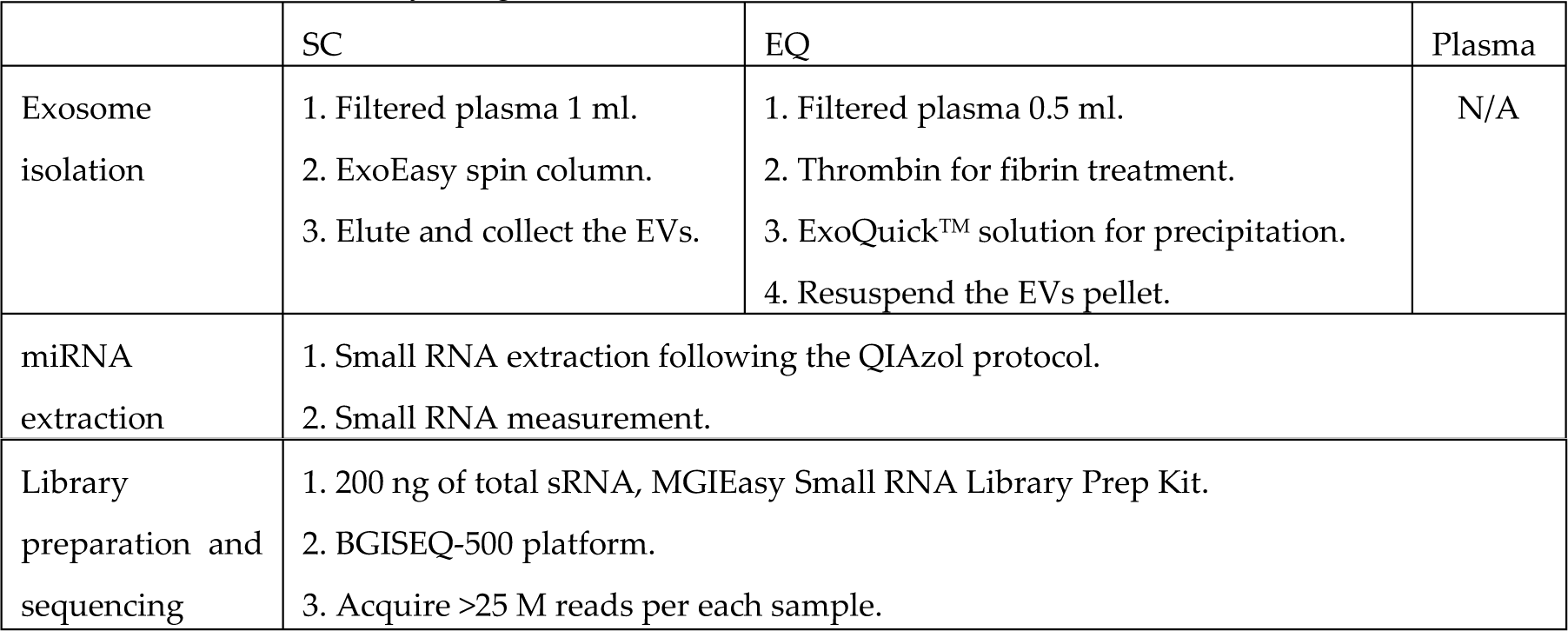
Workflow of the study design.

|  | SC | EQ | Plasma |
| --- | --- | --- | --- |
| Exosome isolation | 1. Filtered plasma 1 ml.<br>2. ExoEasy spin column.<br>3. Elute and collect the EVs. | 1. Filtered plasma 0.5 ml.<br>2. Thrombin for fibrin treatment.<br>3. ExoQuick™ solution for precipitation.<br>4. Resuspend the EVs pellet. | N/A |
| miRNA extraction | 1. Small RNA extraction following the QIAzol protocol.<br>2. Small RNA measurement. |  |  |

|  |  |
| --- | --- |
| Library preparation and sequencing | 1. 200 ng of total sRNA, MGIEasy Small RNA Library Prep Kit.<br>2. BGISEQ-500 platform.<br>3. Acquire >25 M reads per each sample. |

**Figure 1.**
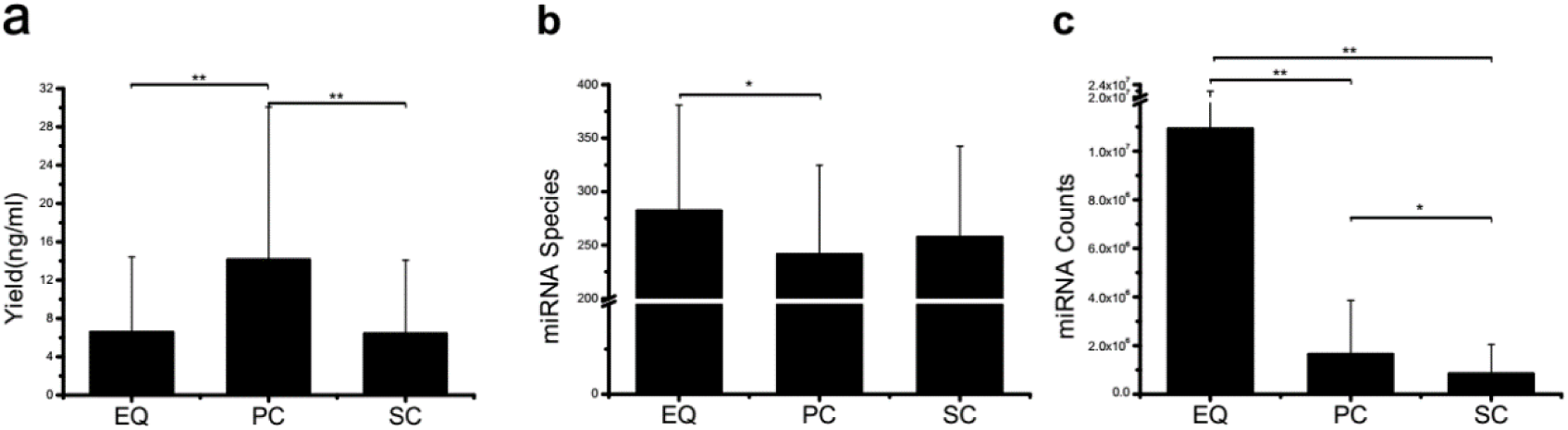
Small RNA yield, miRNA species and total counts from three exosome isolation methods. **a**, small RNA yield efficiency by three methods, average of 46 total production divided by volume of primary plasma, no significant difference was found. **b**, identified miRNA species by three methods, average of miRNA species from 46 samples, EQ exhibited significant more miRNA number than PC. **c**, miRNA total counts by three methods, average of total miRNA counts from 46 samples, EQ presented significantly higher number than other two methods. * presents p < 0.05 in Student-T test. ** presents p < 0.01 in Student-T test.

### 2.2 There are no fundamental differences between SC and EQ in exRNA yield while EQ exhibits better data quality

We preliminarily compared the three datasets, i.e. EQ, SC, and PC, from 46 samples by yield efficiency (small RNA production divided by the volume of plasma), aligned known miRNA from miRBase and miRNA total counts (Figure 1). The total yields of 138 (46X3) samples from EQ, SC, and PC were measured and presented in terms of yield per volume of original plasma (Supplementary Table 2). The average yield of PC is significantly larger than EQ and SC (p < 0.01) and there is no difference between the yield of EQ and SC, even though the intergroup differences of three groups are severe. (Figure 1a). It is indicated that plasma samples may lose great amount of sRNA in both EQ and SC exosome isolation procedures. The individual variation can also be observed in miRNA species (Figure 1b). The maximum of miRNA species is observed on the EQ group, and it is notably larger than that of the PC group (p < 0.05). It is also noticed that no significant difference can be found either between EQ and SC or between PC and SC. However, after we aligned the sRNA reads to miRbase (http://www.mirbase.org/ftp.shtml), the total number of aligned EQ miRNA counts is significant larger (p <0.01) than that of PC and SC miRNA counts (Figure 1c). Meanwhile, the number of PC miRNA counts is larger (p<0.05) than that of SC group. Therefore, it can be concluded that although no fundamental differences of sRNA yield efficiency and miRNA species can be found among EQ, PC and SC datasets, EQ data presents remarkably larger miRNA counts and better data quality. Furthermore, SC data doesn’t show obvious advantages compared to the data prepared by employing PC method in terms of plasma miRNA.

### 2.3 Quality control of the small RNA-seq library preparation and sequencing

As the small RNA in plasma exosome is in small quantity and constantly changing, the small RNA-seq data could be affected by the batch effect in small RNA extraction significantly. In order to evaluate the bias of batch variances, we added HBRR standard samples into each sequencing lane. It is observed that the number of total reads and filtered reads are in a stable range for the HBRR sample, while HBRR-1 appears to have larger ratio of total reads over filtered reads which indicates a better data quality (Figure 2a). To investigate the enrichment of miRNA in different batches, the sRNA length distribution of HBRR samples is presented (Figure 2b). Obviously, most of the reads are within the range of 18 – 25 nt which is consistent with the miRNA size. It indicates the small RNA enrichment in the library construction from different batches are identical. Besides, we compared the miRNA accumulation and the spearman correlation between HBRR samples (Figure 2c, d). Hierarchical clustering analysis shows that the patterns of miRNA expressions of HBRR-1/3/4/5/6 samples are analogous to each other, and HBRR-12 shows that the patterns are slightly changed especially in highly expressed miRNAs (Figure 2c). Yet, the HBRR-12 highly expressed miRNAs are the same with other HBRR samples. These differences can also be observed in pairwise correlation matrix (Figure 2d). The spearman correlation coefficients between HBRR samples are all over 0.85, while the correlation coefficients of HBRR-3 and HBRR-4 are over 0.95 (Figure 2d). For the miRNAs that can be aligned to miRbase, more than 1000 miRNAs are identified in each HBRR sample (Figure 2e). Based on the aforementioned results, it can be concluded that the batch effect is negligible and the library preparation and sequencing platform is eligible to our isolation method comparison.

**Figure 2.**
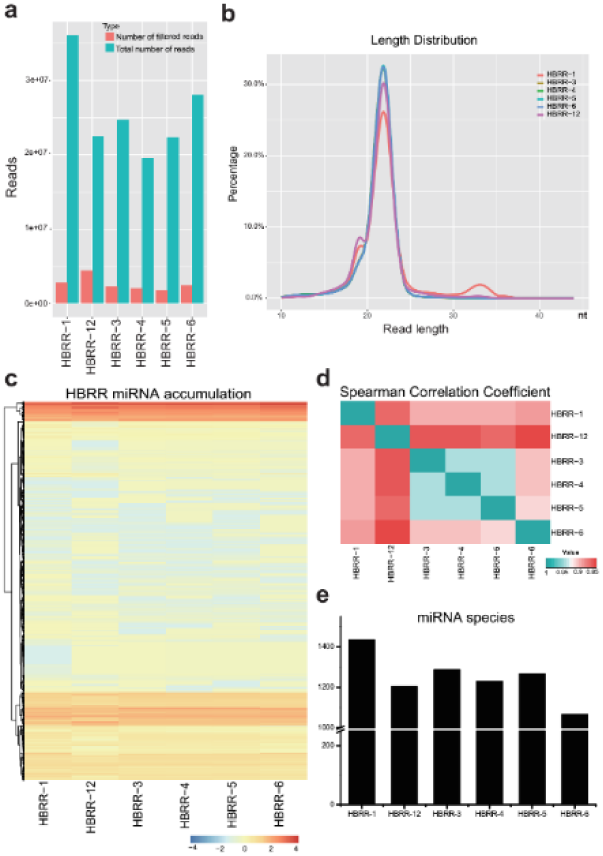
HBRR sRNA data exhibits insignificant sequencing batch effects. **a**, Total reads and filtered reads. **b**, Read length distribution of six HBRR samples. **c**, Heatmap of miRNA accumulation of six HBRR samples. We hierarchically clustered miRNAs based on their miRNA expression (log10). **d**, Pairwise correlation matrix between HBRR samples. Spearman correlation coefficient was calculated and presented as color legend. **e**, Bar chart shows miRNA number identified in each HBRR dataset.

### 2.4 EQ method showed more advantages in terms of data quality

HBRR results exhibit that the batch effect is not a crucial interfering factor in determining different sRNA patterns from EQ, PC and SC datasets, we compared the data quality and known miRNA expression of EQ, PC and SC samples (Figure 3). Firstly, the distribution of small RNA size was plotted from 46 samples of three groups (Figure 3a). EQ samples exhibit high enrichment in 18 – 25 nt range which suggests good sRNA data quality while both SC and PC samples show high percentage of total reads in the < 18 nt range which indicates that small RNA might have some extent of degradation in SC and PC samples (Figure 3a). This assumption of sRNA degradation can also be supported by the aligned miRNA and unmapped reads (Figure 3b). For the same sample, SC samples present the lowest mature miRNA ratio and the highest unmapped ratio, while EQ samples show the highest mature miRNA ratio and the lowest unmapped ratio (Figure 3b). It indicates EQ group has better data quality comparing with the SC and PC groups. Among all detected miRNAs, 695 common miRNAs are found in all three groups (Figure 3c). Meanwhile, 81 miRNAs are only shared by SC samples and EQ samples, 48 miRNAs are only shared by PC samples and EQ samples, 18 miRNAs are only shared by PC samples and SC samples. Additionally, 85, 131, 48 unique miRNAs are observed in SC, EQ and PC samples, respectively (Figure 3c). Among all unique miRNAs, expression of many miRNAs cannot be detected in most of samples (Supplementary Table 3). To illustrate the effect caused by each exosomal isolation method, the pairwise correlation of miRNA profiles for each sample is presented (Figure 3d, e&f). The correlation (R2) values are 0.982 between EQ and PC, 0.954 between EQ and SC, and 0.959 between SC and PC (Figure 3d, e&f), indicating that the sRNA profile of EQ samples is strongly correlated to that of PC samples (Figure 3d). In addition, high correlation values of EQ-SC and SC-PC (>0.95) suggest that all three methods could produce similar data profiles from the same sample.

**Figure 3.**
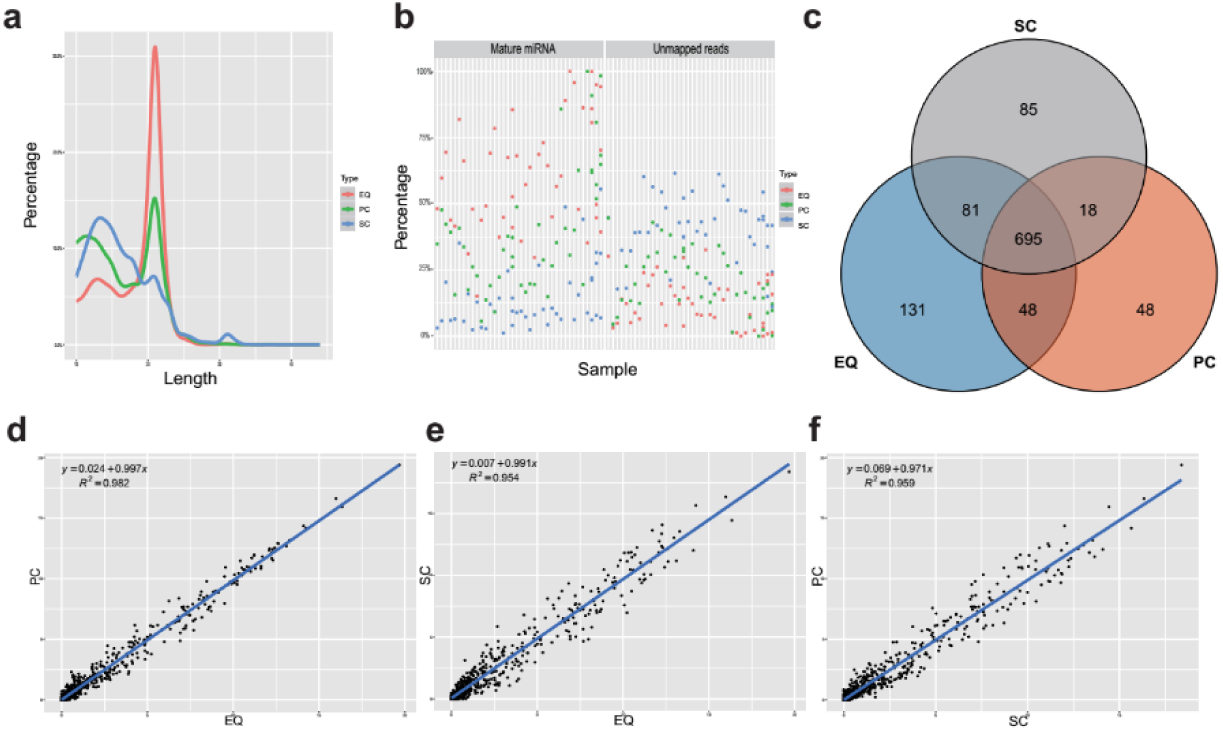
Analysis of the differences in exosomal miRNA profiles among three isolation methods. **a**, Average read length distribution by three methods. **b**, Aligned miRNA reads and unmapped reads of 46 samples by three methods. **c**, Venn diagram presents all identified miRNAs that are common or unique in the three method datasets. Most (695) miRNAs were common to the three sets. **d, e, f**, Scatter plots reveal correlations between EQ and PC (d), EQ and SC (e), SC and PC (f). Each scatter plot represents relative expression of miRNA profiles obtained by two methods.

To further study whether EQ, SC, PC samples could be distinguished by the miRNA expression profiles, principal component analysis (PCA) was performed based on 100 most abundant miRNAs (Figure 4). For EQ and PC samples, 41 out of 46 samples are clustered into Group 2, while 33 out of 46 samples of SC samples are clustered into Group 3. Thus, it indicates that the miRNA profile of SC method can be distinguished from that of EQ and PC methods. Combining the facts that sRNA extracted using PC method contains both exosomal sRNA and plasmatic cell-free sRNA, and exosomal sRNA is better protected by lipid bilayer from RNases, exosomal sRNA profile should have higher miRNA abundance and be included by plasma sRNA profile. Therefore, it can be concluded that EQ method can produce better sRNA data profile than SC method.

**Figure 4.**
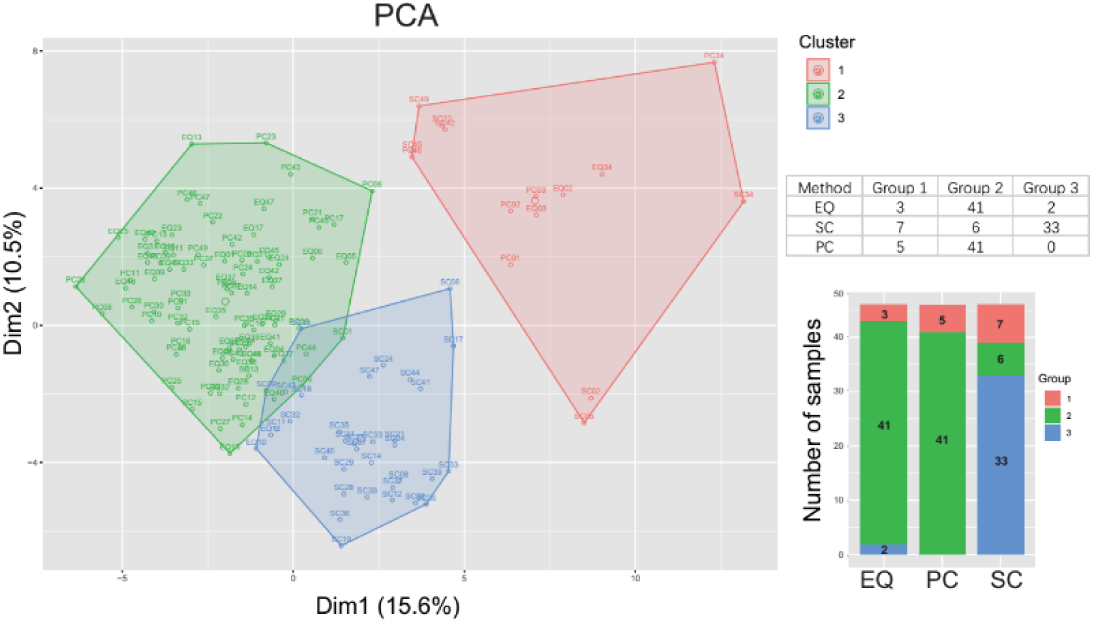
Principal components analysis (PCA) plot of all sample datasets based on their miRNA expression with the colors represent the three clustered groups (left panel). Sample names are presented on top of each dot. Table and bar chart on the right shows the number of samples from three methods that are clustered to each group.

## 3. Discussion

In this study, we compared sRNA sequencing results using two exosome isolation kits named ExoQuick™ exosome precipitation solution and exoEasy spin column. Meanwhile, a blank control group without exosome isolation steps was also added into the comparison. Although there have been several studies comparing different small RNASeq library preparation methods for plasma exosomes, this study is specially focused on the sRNA profile consistency and reproducibility of different exosomal isolation kits. We found that small RNA-seq data acquired by EQ method has a better quality than that obtained by SC method. SC dataset can also be distinguished by miRNA expression (Figure 4). Though we clustered most of EQ (41 out of 46) and PC (41 out of 46) samples into the same group by miRNA expression profiles, data quality of EQ is obviously better than PC samples (Figure 3a, b, Figure 4). Due to the high concentration of RNases in human blood plasma, the cell-free RNAs are fragile and unstable as well as small RNAs except those which can bind to proteins such as AGO2[13,14]. Bilayer lipid structure of exosomes could protect sRNAs from RNases, so that, in our study, exosomal sRNAs should be one of the major contents of the plasma sRNAs[15]. Thus, it is reasonable that sRNA profiles of most EQ samples can be grouped with PC samples in PCA which shows clear clustering of the three methods (Figure 4).

In order to discover whether EQ, PC and SC samples could be distinguished by the miRNA profiles, the k-means clustering was carried out to classify data generated from one sample by EQ, PC and SC methods. To be consistent with PCA, we used 100 most abundant miRNA as the data for clustering (Figure 5). Unsurprisingly, between EQ and PC samples, 3 and 39 common samples are in the Cluster 1 and 2, respectively. Between EQ and SC samples, 2, 6 and 2 common samples are in the Cluster 1, 2 and 3, respectively (Figure 5a). EQ and PC have much more common samples than EQ and SC which indicates that the data of PC samples is in high similarity to data of EQ samples (Figure 5b). Co-inertia analysis between the miRNA profiles of EQ, PC and SC samples reveals a significant co-variation (Figure 5c, d, e)[16]. The RV coefficients of the three pairwise comparisons show that the profiles of EQ and PC samples are more similar than the other two comparisons (Figure 5c, d, e). Taking together, the intergroup variance greatly influences the sRNA yield of EQ and SC samples, and EQ method could produce better sRNA profiles than SC method.

**Figure 5.**
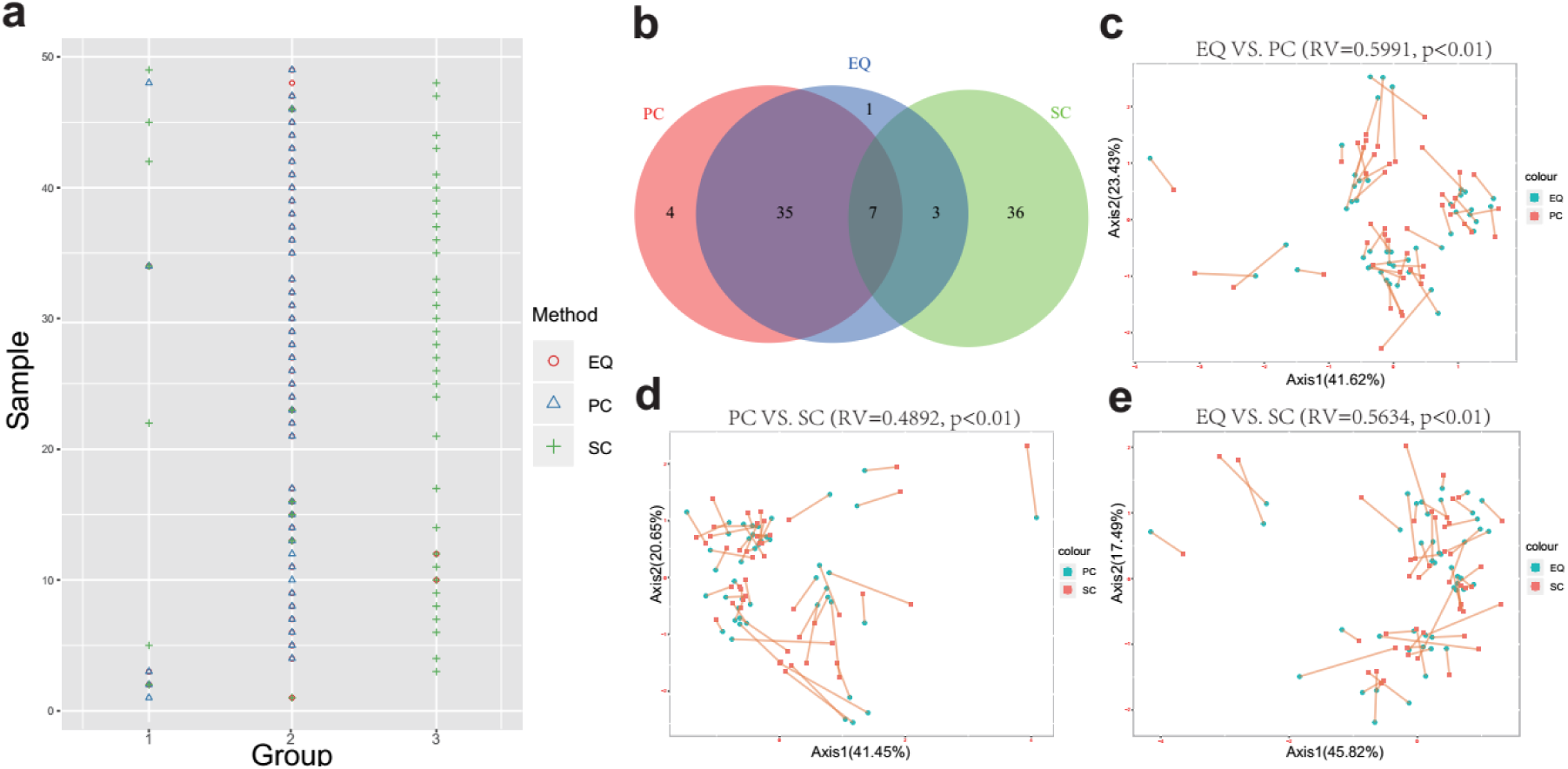
Correlation of EQ, PC and SC profiles. a, Grouping information for each sample by PCA. Colors and shapes present three exosome isolation methods. Y-axis shows the sample number. X-axis shows the grouping information. b, Venn diagram presents EQ, PC and SC samples that are grouped into the same clusters. c, Co-inertia analysis (CIA) of relationships between the EQ and PC miRNA expressional PCA. d, CIA of relationships between the PC and SC miRNA expressional PCA. e, CIA of relationships between the EQ and SC miRNA expressional PCA. c, d, e, p value <0.01 from 99 Monte-Carlo simulations.

In most clinical studies, sample collection usually takes a period of time and collected samples cannot be immediately handled until the quantity of samples is big enough for the study. In order to test whether the storage would affect exosomal sRNA profile, we additionally selected 4 plasma samples and stored them at −80 °C for one, two and three months. Then, we isolated exosome, and carried out sRNA sequencing with the same condition of previous 46 samples. EQ samples exhibit the highest miRNA counts at 1st month in all four samples. Yet, the miRNA counts of EQ samples declined sharply at 2nd month and continued to drop at 3rd month (Figure 6). Except EQ samples, miRNA counts of PC and SC samples doesn’t show a very clear pattern with time. PC samples vary greatly from person to person while the quality (identified reads) of extracted sRNA from SC samples are not good enough for sRNA sequencing (Figure 6). Therefore, in order to get good sRNA data quality, plasma samples should be stored at −80 °C within two months, and use EQ rather than SC for exosome isolation.

**Figure 6.**
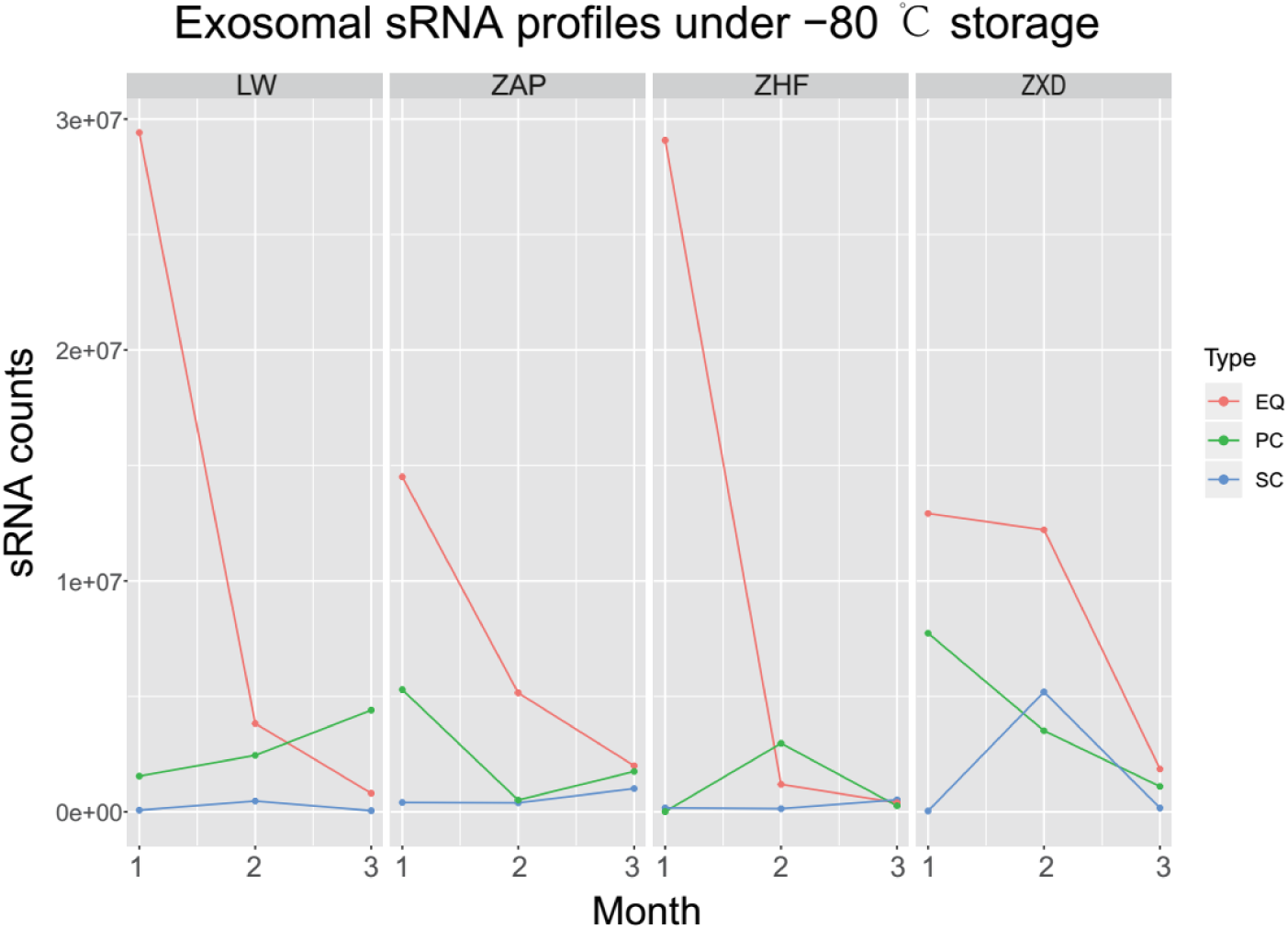
Impact of storage on sRNA data quality. Four samples were collected and stored at −80 ° C condition. Line chart shows the sRNA total counts of four samples after 1 month, 2 months and 3 months storage.

For clinical treatment, exosomes are frequently proposed as therapeutic drug carriers [17]. Six major exRNA cargo types have been discovered in a systematic research of the NIH Extracellular RNA Communication Consortium[18]. Since exRNA cargo and the exosome source are cell type specific, exosomes have been the potential biomarkers for disease diagnosis [19]. Therefore, exosome-based approaches aimed at identifying miRNAs in blood could evaluate the risk of diseases. Additionally, it may also be an alternative strategy that might facilitate the disease diagnosis, especially suitable for diseases of which existing tests are invasive and expensive, such as neurodegenerative diseases[20]. Patients in neurodegenerative risk, are in desperate need of a noninvasive, convenient, and robust method to monitor disease procession. Traditional ultracentrifugation method of exosome isolation does not satisfy the need for high-through and rapidity. Commercially available kits would overcome the obstacles between related research and clinical application, as well as the cost of diagnosis. Recently, sequencing not only exosomal small RNA but also lncRNA to comprehensively discover biomarkers starts to getting more attention[4]. This study compared sRNA sequencing data generated from two exosome isolation kits, it provides a reference for future small RNA studies and biomarker prediction in human plasma exosome.

## 4. Materials and Methods

Single-donor peripheral blood samples were collected from 46 volunteers (No age, gender, health, and BMI limits) under approval from Affiliated Hospital of Jining Medical University IRB (2017-KE-B001) and BGI IRB (No. BGI-IRB16019), and all volunteers were properly consented before samples were collected, all samples had been anonymized for research purposes. The control samples HBRR (Human Brain Reference RNA, Cat. No. AM6050) was supplied by ThermoFisher.

### 4.1 Plasma Separation and EV Isolation

Samples were collected following the fasting blood standard protocol (https://medlineplus.gov/lab-tests/fasting-for-a-blood-test/). Exosome isolation was carried out using two commercially available kits: exoRNeasy Serum/Plasma Maxi Kit (QIAGEN, Hilden, Germany) (SC); ExoQuick Plasma prep and Exosome isolation kit (SBI, Palo Alto, USA) (EQ). For each sample, we used 1 ml plasma for one reaction of SC kit and 0.5 ml plasma for one reaction of EQ kit.

For employing SC method, we prefiltered plasma and mixed the flow-through with 2× binding buffer. Then the solution was added to the exoEasy membrane affinity column and centrifugated for 1 min at 500 x g. The pellets were washed with washing buffer by centrifuging and discarding the flow-through. The pellets were isolated exosome and was re-suspended with nuclease-free PBS.

For employing EQ method, 5 ul of Thrombin [500U/ml] (SBI, Palo Alto, USA) was added into 0.5 ml prefiltered plasma. The mixture was incubated for 5 min and centrifuged for 5 min at 8,000 × g. We added 1/4 volume of ExoQuick Solution to the supernatant and incubated it at 4 °C for 30 min. The mixture was centrifuged at 1,500 × g for 30 min. Finally, Re-suspended the pellets with nuclease-free PBS.

### 4.2 Small RNA extraction, measurements, library preparation and sequencing

The small RNA was extracted according to the manufacturer’s instructions (QIAGEN, Hilden, Germany, Cat No. 217084). The quality of the RNA samples was tested with Agilent 2100 Bioanalyzer RNA Nanochip. The RNA yield was measured by NanoDrop 2000. Small RNA sequencing libraries were constructed using the MGIEasy Small RNA Library Prep Kit (MGI, Shenzhen, China). Input amounts of RNA was at least 200 ng per sample. 25M reads were generated for each sample consequently.

### 4.3 Small RNA sequencing data analysis

After the removal of adaptor sequences and filtering out low-quality reads, the cleaned sRNAs reads were mapped against human reference genome hg19 UCSC and Rfam (version 11.0, http://rfam.xfam.org/) database to discard rRNA-, scRNA-, snoRNA-, snRNA-, and tRNA-associated reads. The remaining reads were aligned and annotated according to precursor and mature miRNAs listed in miRBase (release 21, http://www.mirbase.org/) by BLASTn with a maximum of one nucleotide mismatch per read [21]. The counts of the identified miRNAs were normalized to transcripts per million (TPM). Standardized data was then used for subsequent differential expression analysis. The normalized read counts were then analyzed by the DEseq2 to identify differentially expressed miRNAs [22].

## Supplementary Materials

Supplementary Table 1. Summary of small RNA sequences from 46 libraries.

Supplementary Table 2. Exosomal RNA quantity in the samples isolated using the different methods.

Supplementary Table 3. Expression of small RNAs in EQ, PC and SC samples.

## Author Contributions

YS analyzed data, and drafted the manuscript. HZ, CN designed and performed the experiments. YY collected samples and connected with patients. JY and LL participated in amending the manuscript. XZ and MG provided helpful suggestions and designs during the analysis, and CN was responsible for the overall concept and revising manuscript. All authors read and approved the manuscript.

## Acknowledgments

We thank for the funding support of Science, Technology and Innovation Commission of Shenzhen Municipality under grant No. JCYJ20170412153100794, JSGG20170824152728492 and Shandong Province Medical and Health Science and Technology Development Plan Project (2017WS222).

## Abbreviations

EV: Extracellular Vesicles
IRB: Institutional Review Boards
HBRR: Human Brain Reference RNA
DNBs: DNA nanoballs
RCR: Rolling Circle Replication

## References

1. Bellingham, S.A.; Coleman, B.M.; Hill, A.F. Small RNA deep sequencing reveals a distinct miRNA signature released in exosomes from prion-infected neuronal cells. Nucleic Acids Res 2012, 40, 10937–10949, doi:10.1093/nar/gks832.

2. Alderton, G.K. Diagnosis: Fishing for exosomes. Nat Rev Cancer 2015, 15, 453, doi:10.1038/nrc3990.

3. Valadi, H.; Ekstrom, K.; Bossios, A.; Sjostrand, M.; Lee, J.J.; Lotvall, J.O. Exosome-mediated transfer of mRNAs and microRNAs is a novel mechanism of genetic exchange between cells. Nat Cell Biol 2007, 9, 654–659, doi:10.1038/ncb1596.

4. !!! INVALID CITATION !!!.

5. Rekker, K.; Saare, M.; Roost, A.M.; Kubo, A.L.; Zarovni, N.; Chiesi, A.; Salumets, A.; Peters, M. Comparison of serum exosome isolation methods for microRNA profiling. Clin Biochem 2014, 47, 135–138, doi:10.1016/j.clinbiochem.2013.10.020.

6. Gonzales, P.A.; Pisitkun, T.; Hoffert, J.D.; Tchapyjnikov, D.; Star, R.A.; Kleta, R.; Wang, N.S.; Knepper, M.A. Large-scale proteomics and phosphoproteomics of urinary exosomes. J Am Soc Nephrol 2009, 20, 363–379, doi:10.1681/ASN.2008040406.

7. Taylor, D.D.; Zacharias, W.; Gercel-Taylor, C. Exosome isolation for proteomic analyses and RNA profiling. Methods Mol Biol 2011, 728, 235–246, doi:10.1007/978-1-61779-068-3_15.

8. !!! INVALID CITATION !!! [6,7].

9. Kamm, R.C.; Smith, A.G. Nucleic acid concentrations in normal human plasma. Clin Chem 1972, 18, 519–522.

10. !!! INVALID CITATION !!! [9,10].

11. Tang, Y.T.; Huang, Y.Y.; Zheng, L.; Qin, S.H.; Xu, X.P.; An, T.X.; Xu, Y.; Wu, Y.S.; Hu, X.M.; Ping, B.H., et al. Comparison of isolation methods of exosomes and exosomal RNA from cell culture medium and serum. Int J Mol Med 2017, 40, 834–844, doi:10.3892/ijmm.2017.3080.

12. Lewis, G.D.; Metcalf, T.G. Polyethylene glycol precipitation for recovery of pathogenic viruses, including hepatitis A virus and human rotavirus, from oyster, water, and sediment samples. Appl Environ Microbiol 1988, 54, 1983–1988.

13. Vickers, K.C.; Palmisano, B.T.; Shoucri, B.M.; Shamburek, R.D.; Remaley, A.T. MicroRNAs are transported in plasma and delivered to recipient cells by high-density lipoproteins. Nat Cell Biol 2011, 13, 423–433, doi:10.1038/ncb2210.

14. Chendrimada, T.P.; Gregory, R.I.; Kumaraswamy, E.; Norman, J.; Cooch, N.; Nishikura, K.; Shiekhattar, R. TRBP recruits the Dicer complex to Ago2 for microRNA processing and gene silencing. Nature 2005, 436, 740–744, doi:10.1038/nature03868.

15. Huang, X.; Yuan, T.; Tschannen, M.; Sun, Z.; Jacob, H.; Du, M.; Liang, M.; Dittmar, R.L.; Liu, Y.; Liang, M., et al. Characterization of human plasma-derived exosomal RNAs by deep sequencing. BMC Genomics 2013, 14, 319, doi:10.1186/1471-2164-14-319.

16. Zhao, L.; Zhang, F.; Ding, X.; Wu, G.; Lam, Y.Y.; Wang, X.; Fu, H.; Xue, X.; Lu, C.; Ma, J., et al. Gut bacteria selectively promoted by dietary fibers alleviate type 2 diabetes. Science 2018, 359, 1151–1156, doi:10.1126/science.aao5774.

17. Lin, J.; Li, J.; Huang, B.; Liu, J.; Chen, X.; Chen, X.M.; Xu, Y.M.; Huang, L.F.; Wang, X.Z. Exosomes: novel biomarkers for clinical diagnosis. ScientificWorldJournal 2015, 2015, 657086, doi:10.1155/2015/657086.

18. !!! INVALID CITATION !!! [14].

19. Thery, C.; Zitvogel, L.; Amigorena, S. Exosomes: composition, biogenesis and function. Nat Rev Immunol 2002, 2, 569–579, doi:10.1038/nri855.

20. Abdel-Haq, H. Blood exosomes as a tool for monitoring treatment efficacy and progression of neurodegenerative diseases. Neural Regen Res 2019, 14, 72–74, doi:10.4103/1673-5374.243709.

21. Altschul, S.F.; Gish, W.; Miller, W.; Myers, E.W.; Lipman, D.J. Basic local alignment search tool. Journal of Molecular Biology 1990, 215, 403–410, doi:http://dx.doi.org/10.1016/S0022-2836(05)80360-2.

22. Love, M.I.; Huber, W.; Anders, S. Moderated estimation of fold change and dispersion for RNA-seq data with DESeq2. Genome Biol 2014, 15, 550, doi:10.1186/s13059-014-0550-8.

